# Hierarchical Regulation of Autophagy During Adipocyte Differentiation

**DOI:** 10.1101/2021.01.06.425505

**Authors:** Mahmoud Ahmed, Trang Huyen Lai, Trang Minh Pham, Sahib Zada, Omar Elashkar, Jin Seok Hwang, Deok Ryong Kim

**Affiliations:** Department of Biochemistry and Convergence Medical Sciences and Institute of Health Sciences, Gyeongsang National University School of Medicine, Jinju, South Korea

**Keywords:** transcription-factors, autophagy, differentiation, adipocyte, hierarchical-regulation

## Abstract

We previously showed that some adipogenic transcription factors such as CEBPB and PPARG directly and indirectly regulate autophagy gene expression in adipogenesis. The order and the effect of these events are undetermined. In this study, we modeled the gene expression, DNA-binding of transcriptional regulators, and histone modifications during adipocyte differentiation and evaluated the effect of the regulators on gene expression in terms of direction and magnitude. Then, we identified the overlap of the transcription factors and co-factors binding sites and targets. Finally, we built a chromatin states model based on the histone marks and studied their relation with the factors’ binding. Adipogenic factors differentially regulated autophagy genes as part of the differentiation program. Co-regulators associated with specific transcription factors and preceded them to the regulatory regions. Transcription factors differed in the binding time and location, and their effect on expression was either localized or long-lasting. Adipogenic factors disproportionately targeted genes coding for autophagy-specific transcription factors. To sum, a hierarchical arrangement between adipogenic transcription factors and co-factors drives the regulation of autophagy during adipocyte differentiation.

## 1 Introduction

Previous studies suggested one-to-one interactions between adipogenic transcription factors and autophagy. CEBPB transactivates *Atg4b*, a key protein in the autophagy machinery [1]. The activation of autophagy through this pathway relieves the repression of adipogenic activators such as PPARG. FOXO1, a transcription factor with several autophagy targets, was suggested to the repress *Pparg* gene in the presence of insulin sensitizers [2]. This repression is likely to be lifted in early adipogenesis. A previous study from our laboratory showed that autophagy gene products are regulated as part of the transcription program of adipogenesis [3]. This regulation is achieved through adipogenic transcription factors PPARG and CEBPB either directly or indirectly through autophagy specific factors. The magnitude and the ordering of this regulation remain to be investigated.

Here, we used gene expression and DNA-binding data to model the transcription factor and co-factors binding events during differentiation and their effect on autophagy genes. We used histone modification data to correlate these events with chromatin states. A hierarchical arrangement of known adipogenic transcription factors and co-factors emerged in the regulation of autophagy during adipogenesis. We evaluated the spatial and temporal aspects of this arrangement. These included the factors’ contributions to gene expression, the dependency between the regulators, the reliance on chromatin states, and the type of binding targets.

## 2 Methods

### 2.1 Expression & binding data

We collected two datasets of RNA-seq and ChIP-seq on 3T3-L1 pre-adipocytes, which were induced to differentiate using 3-isobutyl-1-methylxanthine, dexamethasone, and insulin (MDI) and sampled at different time points (Table 1, 2, & 3) [4]. We curated the samples’ metadata using a unified language across the studies and processed the raw data using standard pipelines. The processed gene expression data were made available as a Bioconductor data package (curatedAdipoRNA). The data are presented as gene counts at different time points (0 to 240 hr). The processed DNA-binding data of transcription factors, co-factors, and histone modifications were made available as a similar package (curatedAdipoChIP). Data are presented in this package as the reads count in a consensus peak set. Moreover, we provided links to the identified peaks as well as the signal tracks. The packages document the pre-processing and processing pipelines.

**Table 1:** MDI-induced 3T3-L1 gene expression data by RNA-seq.

| GEO ID | N | Time (hr) | Ref. |
| --- | --- | --- | --- |
| GSE100056 | 2 | 24 | [10] |
| GSE104508 | 3 | 192 | [11] |
| GSE35724 | 3 | 192 | [12] |
| GSE50612 | 4 | 0/144 | [13] |
| GSE50934 | 6 | 0/168 | [14] |
| GSE53244 | 3 | 0/48/240 | [15] |
| GSE57415 | 4 | 0/4 | [16] |
| GSE60745 | 12 | 0/24/48 | [17] |
| GSE64757 | 6 | 168 | [18] |
| GSE75639 | 3 | 0/48/168 | [19] |
| GSE84410 | 5 | 0/4/48 | [20] |
| GSE87113 | 5 | 0/2/4/48/168 | [21] |
| GSE89621 | 3 | 240 | [22] |
| GSE95029 | 8 | 0/48/144/192 | [23] |
| GSE95533 | 10 | 4/0/24/48/168 | [24] |
| GSE96764 | 6 | 0/2/4 | [25] |

**Table 2:**
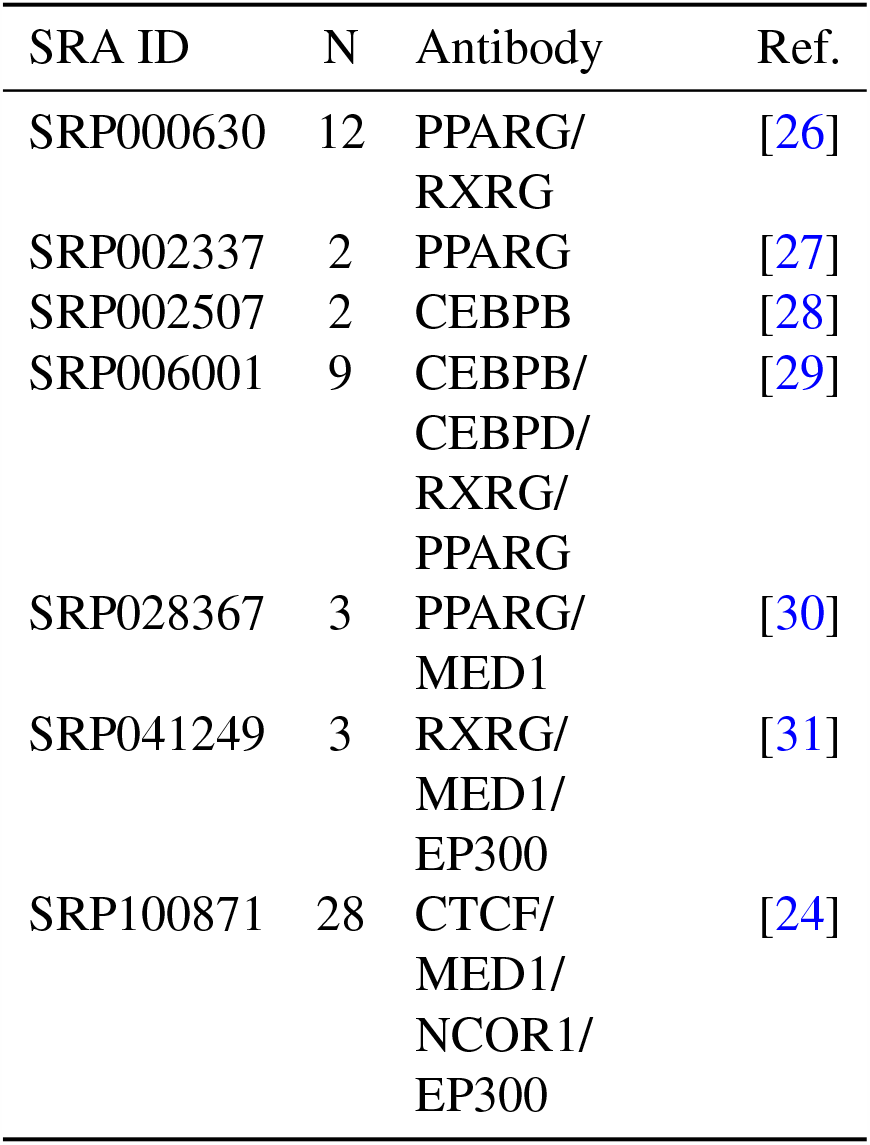
Transcription factors binding data.

**Table 3:** Histone modification data.

| SRA ID | N | Antibody | Ref. |
| --- | --- | --- | --- |
| SRP002337 | 11 | H3K4me3/<br>H3K27me3/<br>H3K36me3/<br>H3K4me2/<br>H3K4me1/<br>H3K27ac | [27] |
| SRP041249 | 6 | H3K27ac/<br>H3K4me1/<br>H3K4me2 | [31] |
| SRP064188 | 3 | H3K27me3/<br>H3K9me3 | [32] |
| SRP078506 | 6 | H3K4me3 | [20] |
| SRP100871 | 6 | H3K27ac/<br>H3K4me1/<br>H3K4me2 | [24] |

We obtained two gene expression datasets of *Cebpb* (RNA-seq) or *Pparg*-knockdown (microarrays) from a matching MDI-induced 3T3-L1 pre-adipocytes time-course experiments (Table 4). Gene counts and probe intensities were downloaded using GEOquery and used to quantify the gene expression from RNA-seq and microarray data, respectively [5].

**Table 4:** Perturbed MDI-induced 3T3-L1 gene expression data by RNA-seq.

| GEO ID | N | KD | Ref. |
| --- | --- | --- | --- |
| GSE57415 | 8 | Cebpb | [16] |
| GSE12929 | 18 | Pparg | [33] |

### 2.2 Mouse genome annotations

Gene ontology (GO) terms in mouse biological processes were used to identify the gene products relevant to autophagy and lipogenesis [6]. The Bioconductor package org.Mm.eg.db was used to access the GO annotations [7]. The gene accessor IDs were mapped between gene symbols and Entrez IDs using TxDb.Mmusculus.UCSC.mm10.knownGene [8]. The same package was used to extract gene coordinates in the mouse genome. Finally, the GO terms for molecular functions terms were used to identify the transcription binding targets’ functional categories. GO.db was used to access these terms [9].

### 2.3 Differential gene expression

RNA-seq reads were aligned to the mm10 mouse genome and counted in known genes using HISAT2, and featureCount [34, 35]. Gene counts were filtered, normalized, transformed, and subjected to batch effects removal. Microarrays probe intensities were filtered and collapsed to corresponding known genes, normalized, and transformed. To identify gene expression changes over time or in response to transcription factors genes knockdown, we applied differential gene expression analysis using DESeq2, or LIMMA [36, 37]. Briefly, the gene counts or the probe intensities were compared between conditions (#hr vs. 0 hr or knockdown vs. control). Fold-change and p-value for every gene in each comparison were calculated. False-discovery rate (FDR) was used to adjust for multiple testing.

### 2.4 Binding peaks analysis

ChIP-seq reads were aligned to the mm10 mouse genome using BOWTIE2 [38]. Binding peaks were identified using MACS2 with the annotation file of the same genome [39]. Peaks were annotated and assigned to the nearest gene using ChIPSeeker [40]. The numbers of binding sites and targets were calculated in each sample. When more than one sample was available for a given factor, only replicated binding sites or targets were included. The intersections of binding sites and targets among the samples were calculated and visualized using ggupset.

### 2.5 Hidden markov chain models

Multi-states hidden Markov chain models of transcription factors and histone modifications in differentiating adipocytes were built using ChromHMM [41]. Briefly, aligned ChIP-seq reads were binarized to 100/200 bp windows over the mm10 mouse genome. Multivariate hidden Markov chains were used to model the factor/marker’s presence or absence in combinatorial and spatial patterns (states). Emission and transition probabilities for the states were calculated to express the probability of each factor/marker being in a given state and the probability of the states transitioning to/from another at different time points. State enrichment over genomic locations and around the transcription start sites was calculated. The R package segmentr (under development) was used to call ChromHMM, capture, and visualize the output.

### 2.6 Gene set enrichment and over-representation

To calculate GO terms’ enrichment scores at different times of differentiation, we ranked all genes by fold-change, performed a walk of the gene set members over the ranked list, and compared it to random walks. The enrichment score is the maximum distance between the gene set and the random walk [42]. ChromHMM calculates the enrichment of states as (C/A)/(B/D) where A is the number of bases in the state, B is the number of bases in external annotation, C is the number of basses in the state, and the annotation and D is the number of bases in the genome. clusterProfiler calculates the over-representation as the number of items in the query and subject groups compared to the groups’ total number [43].

### 2.7 Software & reproducibility

The analysis was conducted in R language and environment for statistical computing and graphics [44]. Several Bioconductor packages were used as data containers and analysis tools [45]. The software environment was packaged as Docker image (https://hub.docker.com/r/bcmslab/adiporeg). The code for reproducing the analysis and generating the figures and tables in this manuscript is released under GPL-3 open source licence (https://github.com/BCMSLab/hierarchical_autophagy_regulation).

## 3 Results

### 3.1 Adipogenic factors regulate autophagy genes during differentiation

To examine the expression of autophagy genes during adipogenesis, we used a dataset of MDI-induced 3T3-L1 pre-adipocytes sampled at different time points and profiled by RNA-Seq. In addition, we used two datasets of a similar time-courses with *Cebpb* or *Pparg* perturbations. We found that preadipocytes responded to MDI induction by changes in gene expression as early as 4 hours and continue for days. The size of the response was reasonably stable during the differentiation and was evenly split (25% at a false-discovery rate (FDR) *<* 0.2) between genes regulated in either direction. The response was strong for adipogenesis and lipogenesis genes. A larger fraction (30% at day 2 and 50% at day 7 at FDR *<* 0.2) of the genes involved in these processes were progressively induced (up-regulated) up until day 7 of differentiation.

The autophagy response to the MDI induction is bi-phasic with an inflection point around day 2 (Figure 1A). The initial response involved the down-regulation of most autophagy genes (*>* 40% at FDR *<* 0.2). This pattern was reversed in the latter days, where many more autophagy genes were upregulated (40% at day 7 at FDR *<* 0.2). At the gene set level, the products in the gene ontology (GO) term “autophagy” were represented in the down-regulated set (normalized enrichment score (NES) *<* −1.3) up to day 2 and in the up-regulated set (NES *>* 0.8) from then onward (Figure 1B). By contrast, the gene products in the GO term “lipid metabolic process” were always represented in the up-regulated (NES *>* 1.2) ranks in the list of genes.

**Figure 1:**
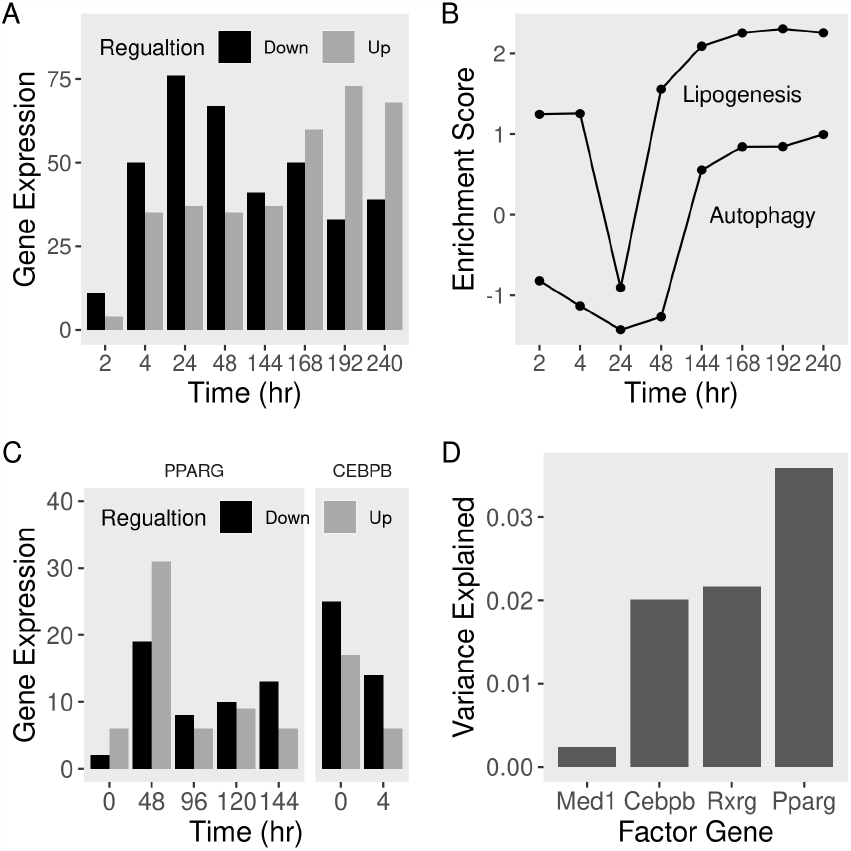
Expression of autophagy gene products during adipocyte differentiation. Read counts were used to quantify the expression of autophagy (and lipogenesis) genes at different times points of differentiation with or without perturbations. Expression was compared to pre-adipocytes (0 hr) to calculate fold-change and p-values. Genes were descendingly ranked by fold-change. A) Number of up-or down-regulated genes in the non-modified course. B) Over-representation of the “autophagy” and “lipogenesis” term members in the top or bottom ranks of the list. C) Number of up-or down-regulated genes with *Pparg*-or *Cepbp*-perturbations. D) Fraction of expression variance explained by the adipogenic regulators’ coding genes.

Adipogenic transcription factors such as CEBPB and PPARG drive autophagy gene expression changes. The expression of the genes coding for those two transcription factors was induced (log_2_ fold-change (FC) *>* 1.75 and 2.5 at day 2 at FDR *<* 0.01, respectively). The knockdown of these factors in pre-adipocytes produced wide dysregulation of autophagy genes (Figure 1C). *Pparg*-knockdown resulted in the up-regulation of 5 to 30 (FDR *<* 0.2) autophagy genes during the first 48 hours of MDI-induction. More than ten autophagy genes were down-regulated (FDR *<* 0.2) by the factor knockdown in the later stages of differentiation. *Cebpb*-knockdown, on the other hand, resulted in the down-regulation (25 to 15 genes at FDR *<* 0.2) of autophagy genes in preadipocytes four hours after MDI induction. Overall, PPARG explains more of the variance in autophagy gene expression (3.5%) than CEBPB (2%) or co-factors (Figure 1D).

### 3.2 Autophagy subtypes exhibit stage-dependent activation

In agreement with the previous literature, we observed the induction of CEBPB and PPARG in early and intermediate adipogenesis. However, we could identify binding sites for both factors at all times points and in preadipocytes. The targets of CEBPB seem to be regulated for a brief period of time that coincided with the induction of the *Cebpb* expression and doesn’t last for long (Figure 2B). By contrast, PPARG binding induced its targets’ expression, and the induction lasted till the end of the experiment (Figure 2A).

**Figure 2:**
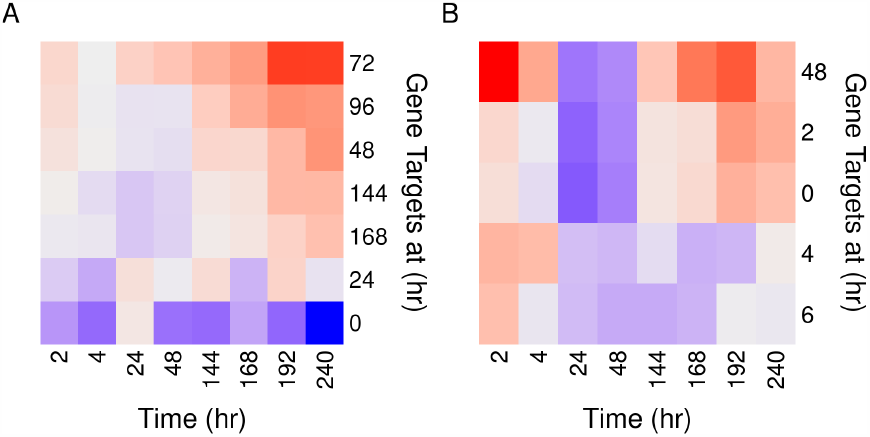
Expression of adipogenic transcription factor targets in the course of differentiation. Binding peaks of A) PPARG or B) CEBPB were used to identify autophagy gene targets at every time point of adipocyte differentiation, rows. Read counts were used to quantify the expression of autophagy genes at all times, columns. Expression was compared to pre-adipocytes (0 hr) to calculate fold-change and p-values. Median fold-change of each target set (blue, low & red, high).

To further explore the effect of the factor binding on autophagy, w calculated the enrichment scores of several autophagy-related terms at the different time points of differentiation (Figure 3). The term “negative regulation of autophagy” was enriched in the downregulated genes in the first two days of differentiation. The was reversed after 48 hours along. Besides, the positive regulation term was later enriched in up-regulated genes. Organelle-specific autophagy terms were enriched after the same time point (48 hr). Terms in the autophagy subtypes that related to the same organelles were also enriched in the up-regulated genes in late-adipogenesis. Together, the biphasic response of autophagy to the MDI-induction was significant in terms of the number of regulated genes and at the gene set level. In particular, selective and organelle-specific forms of autophagy were activated in late-adipogenesis.

**Figure 3:**
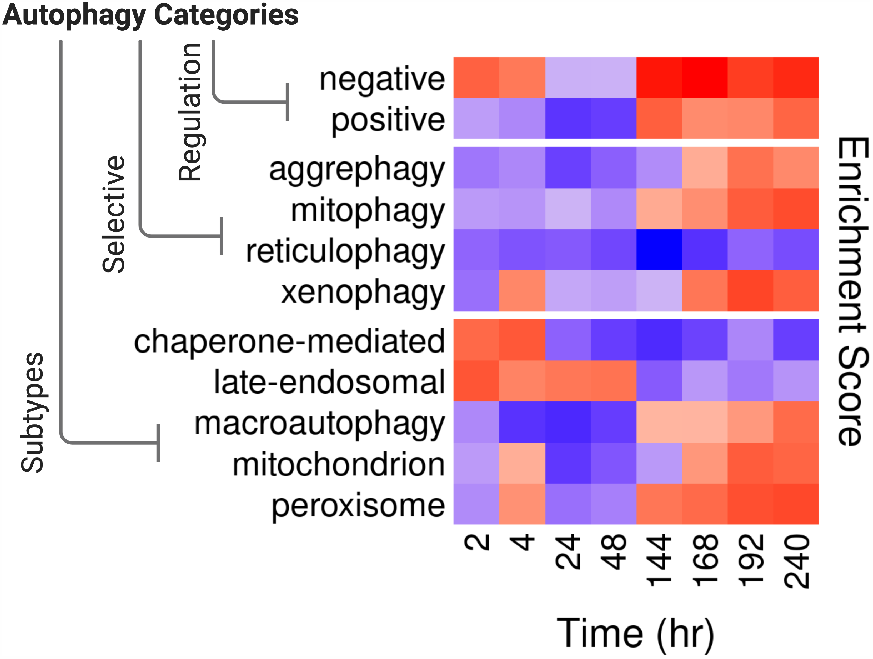
Enrichment of autophagy regulation and subtypes terms at different time points of differentiation. Read counts were used to quantify the expression of autophagy genes at different times points of differentiation. Expression was compared to pre-adipocytes (0 hr) to calculate fold-change and descendingly rank the genes. Enrichment of the “regulation,” “selective,” and “subtypes” terms members in the top or bottom ranks of the list (blue, low & red, high).

### 3.3 Co-regulators are recruited to ubiquitously bound autophagy gene regions and redistribute over time

We further explored the combined binding of key adipogenic factors and co-factor at genome-scale using a ten-states model of the chromatin at the early and later stages of differentiation. During the early stages of differentiation (4 hours), regions of the chromatin fell into one of two categories demarcated by binding patterns (Figure 4A). The first were either devoid of binding proteins (90%), insulated by CTCF (1%), or repressed by NCOR1 (1%) regardless of the presence of other proteins. These areas were generally stable and mainly transitioned to other states of the same category. The active areas were ubiquitously bound to multiple proteins, cofactors only, or a specific transcription factor with its known co-factor. CEBPB associated with MED1 (emission probability (EP) = 0.56 and 0.46, respectively) and PPARG associated with RXRG (EP = 0.86 and 0.85, respectively). These regions were likely to make the transition either from similar binding patterns or from areas devoid of factor binding. To sum, regions that are not open for binding remain so. The binding of transcription factors is sometimes associated with insulators or repressors and is mostly accompanied by co-factors.

**Figure 4:**
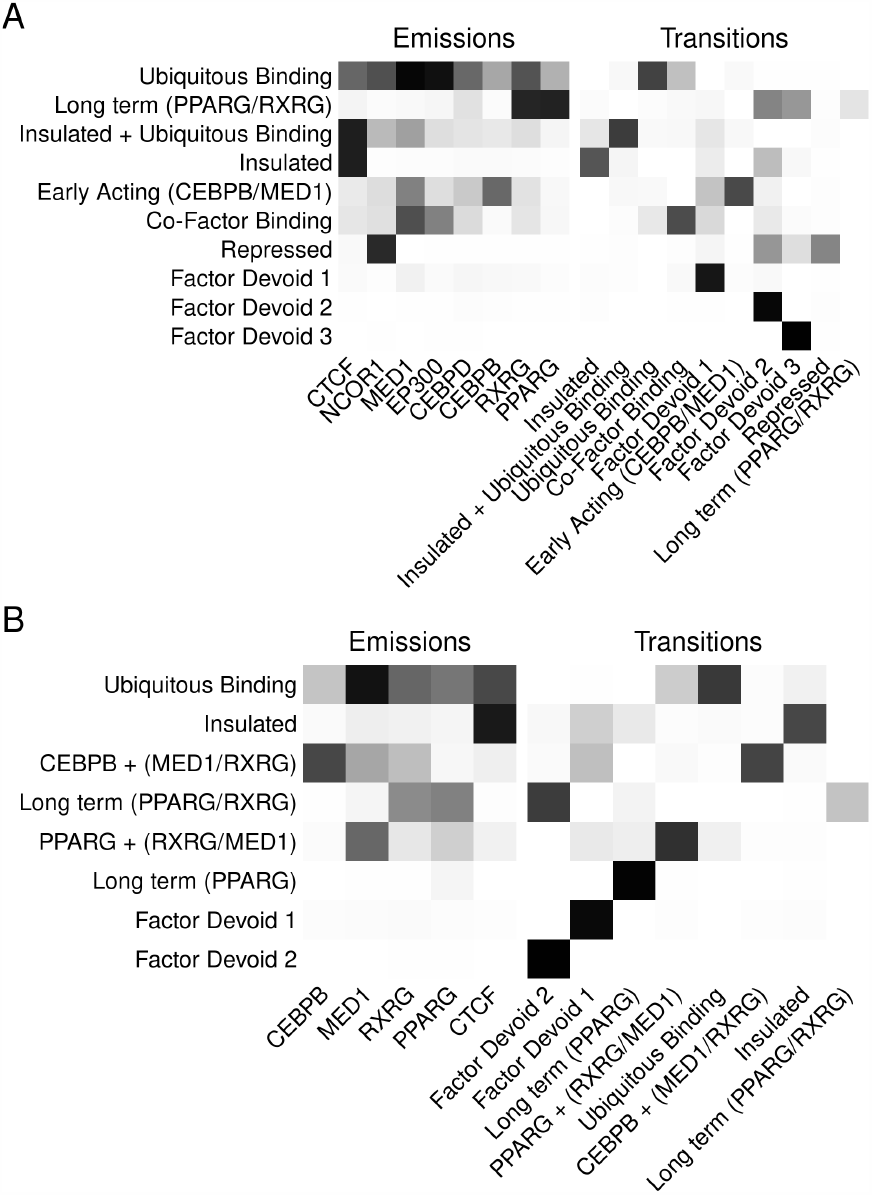
Transcriptional regulators multi-state model in early and full differentiation courses. Binarized binding signals at 100 bp windows were used to indicate the presence or absence of regulators on the chromatin. A multi-variate model of combinations of regulators was built to summarize A) ten states in the early stage or B) eight states in the full course of differentiation. Emission, the probability of each marker being at a given state. Transition, the transitional probability of a given state from/to another. white, low, and black, high probability.

The same patterns of transcription factor and co-factor combinations in an eight-states model emerged at the later stages of differentiation with notable additions (Figure 4B). Repressed regions were more stable and less likely to transition to other states. Transcription factors CEBPB and PPARG associated with more than a single co-factor. In the case of PPARG, the complex of the transcription factors and co-factors (RXRG and MED1) made the transition from the earlier (PPARG + MED1) or PPARG alone states. However, the CEBPB complex made the transition to areas devoid of factors. Possibly, co-factors allow in, or themselves are being recruited by transcription factors to regions with high binding affinities.

In early adipogenesis, significant changes in the states that pertain to insulation, repression, or the binding availability of the chromatin took place. On both autophagy and lipogenic genes, the frequency of the chromatin regions ubiquitously bound to regulatory proteins and co-factors in particular increased (*>* 3 fold) (Figure 5A). This was also accompanied by reduced binding to insulators and repressors (*>* 2 fold). However, in the longer course of differentiation, the more pronounced changes in state frequencies involved the combinations of short and long-acting transcription factors and their association with specific co-factors (Figure 5B). Fewer regions were available for the CEBPB/MED1 complex, and more were available for PPARG either alone or in association with RXRG and MED1.

**Figure 5:**
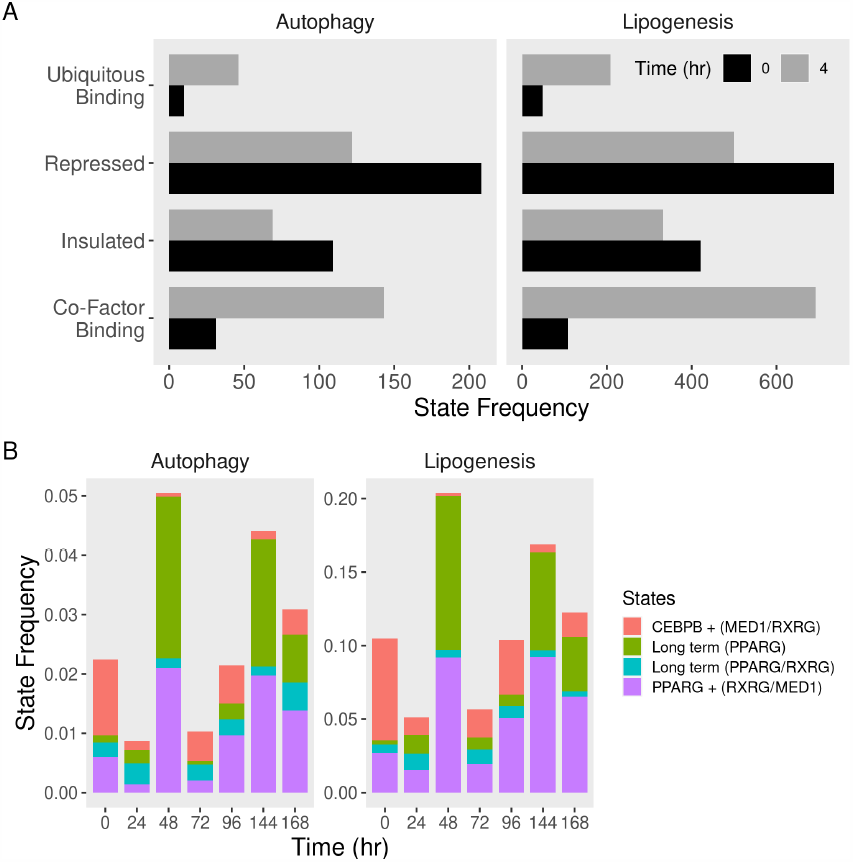
Frequencies of transcriptional states of autophagy gene regions in differentiating adipocytes. Binarized binding signals at 100 bp windows were used to indicate the presence or absence of regulators on the chromatin and build multi-variate models of their combinations during adipogenesis. Autophagy and lipogenesis genomic regions were segmented and labeled by the corresponding states. Frequencies of selected sates were calculated at each time point of the A) early-stage and B) full course of differentiation.

### 3.4 Adipogenic factors are preceded by co-factors on their targets

To examine the co-occurrence of the adipogenic transcription factors on autophagy, we analyzed the intersections of the features they bind to at different time points. PPARG targeted the largest numbers of autophagy genes (Figure 6A). Those targets localized in the later time points. By contrast, CEBPB had the largest number of targets in pre-adipocytes and early after induction with MDI. The downstream targets of PPARG overlapped with those of RXRG, while CEBPB targets overlapped with MED1, especially in early time points. Moreover, co-factors such as MED1 and RXRG and, to a lesser extent CEBPB, accessed their targets independent of the time point. This is confirmed by calculating the fraction of overlap between the PPARG and CEBPB binding with that of the co-factors (Figure 6B). Unlike CEBPB, the fraction on PPARG binding targets with the targets of the co-factors increased over time.

**Figure 6:**
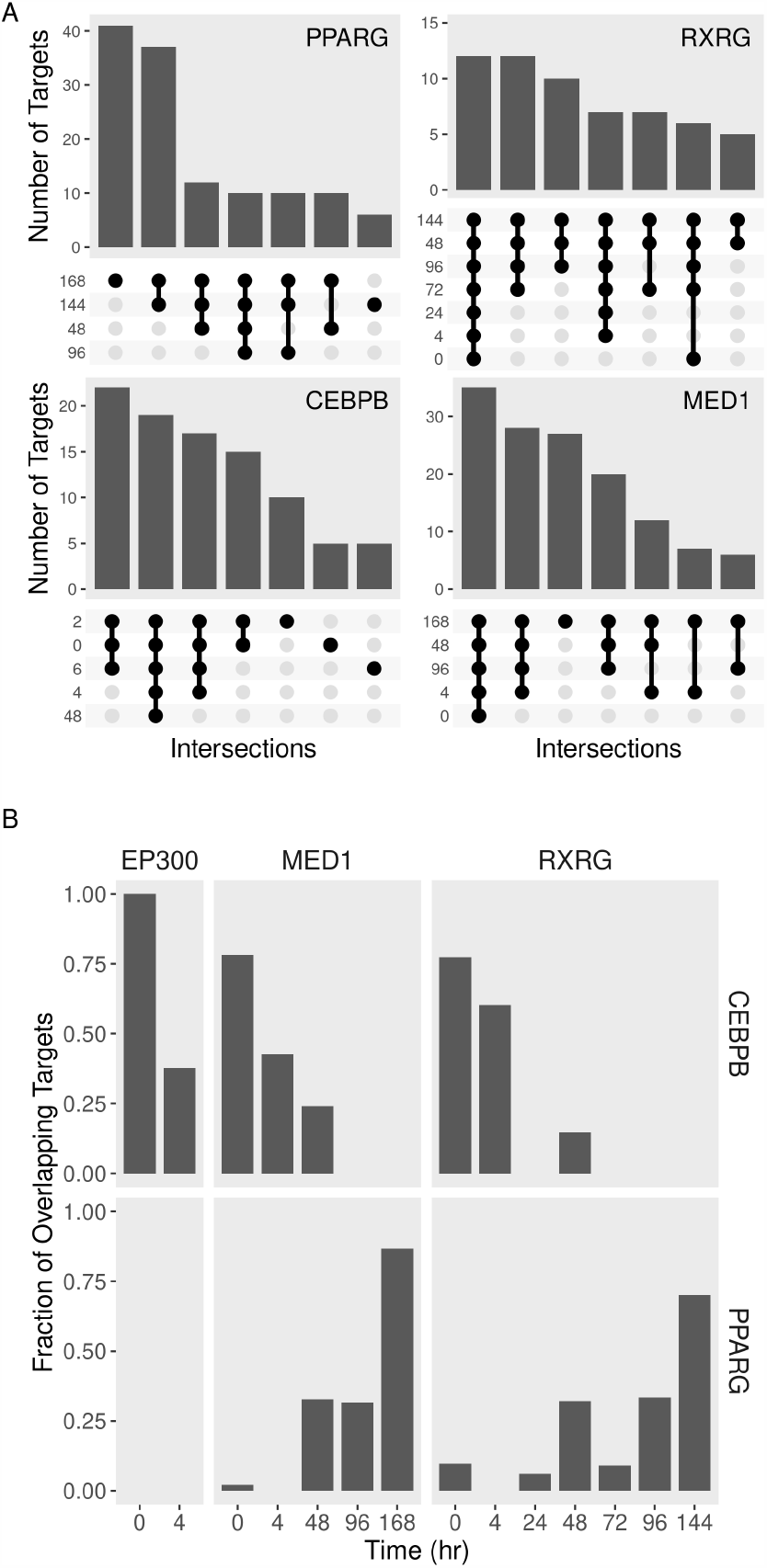
Autophagy gene target of adipogenic regulators and their overlap at different time points. Binding peaks of adipogenic regulators were used to identify gene targets during the course of adipocyte differentiation. A) Numbers of genes in the intersections of the regulators’ targets at different time points. B) Fractions of overlap between transcription factors and co-factors targets at different times.

### 3.5 Co-factors localize to and prime gene enhancers for transcription factors

We then constructed a multi-state chromatin model of histone modifications and examined their co-occurrence with different individual factors and the combinations. Chromatin regions fell into one of two general categories: active or repressed chromatin (Figure 7A). The combination of H3K27ac and H3K4me1 marks the enhancer regions (EP *>* 0.7 and 0.9), while the combination of H3K27ac and H3K4me3 (EP *>* 0.8 and 0.9) marks the active promoters. H3K36me3 marks regions with strong transcriptional activity. The enhancers were further classified into active, weak, or genic depending on the distribution of the histone markers. Active regions mainly transitioned within the same category of enhancers and genic enhancers to robust transcription. The inactive chromatin was annotated by either H3K27me3 (Repressed poly-comb), H3K9me3 (Repeats), or devoid of any markers (Heterochromatin). The only transition from inactive to active regions occurred between weak enhancers and repeats.

**Figure 7:**
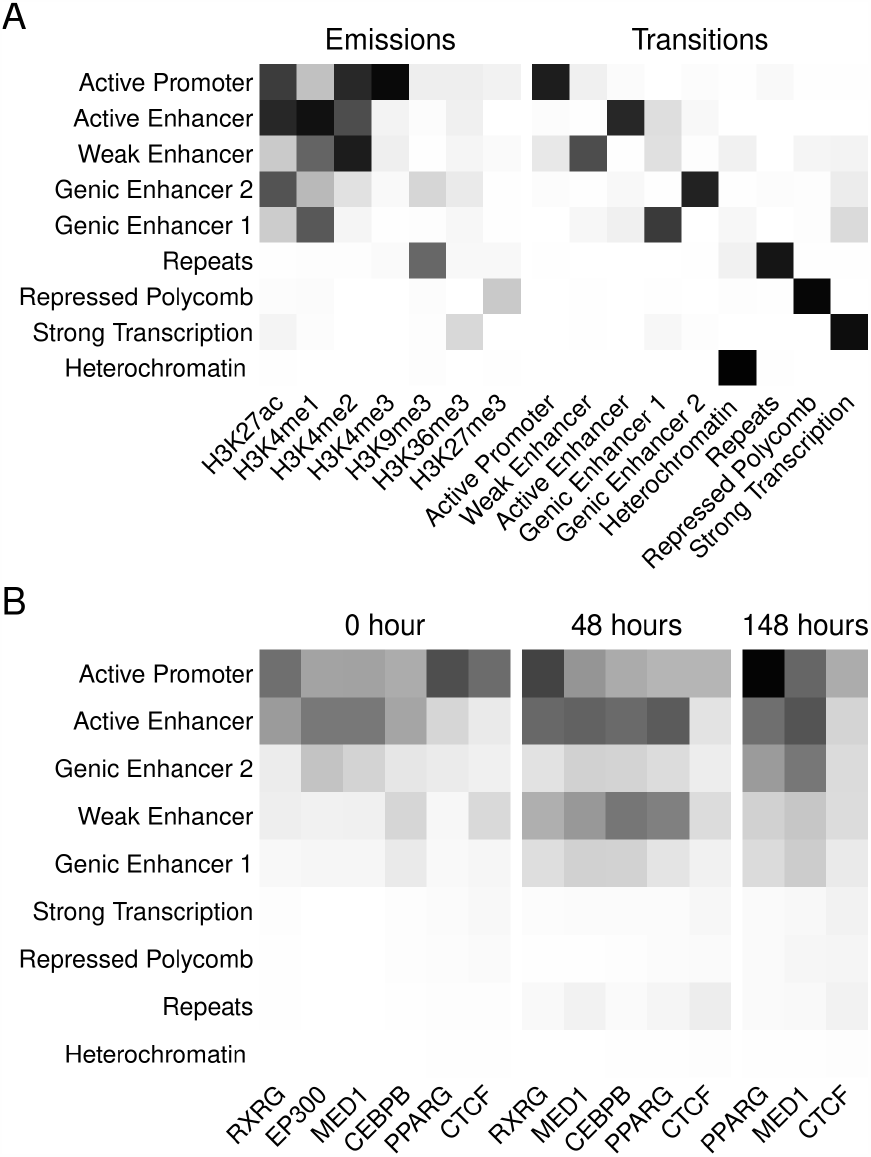
Histone modifications multi-state model and overlap with adipogenic regulators. Binarized binding signals at 200 bp windows were used to indicate the presence or absence of histone marks on the chromatin. A multi-variate model of combinations of marks was built to summarize nine states during the course of differentiation. Binding peaks of adipogenic regulators were used to identify gene targets at the corresponding time points. A) Emission, the probability of each marker being at a given state. Transition, the transitional probability of a given state from/to another. B) Overlap of the regulators binding sites in each chromatin state. white, low, and black, high probability, or overlap.

Autophagy genes regulatory regions such as promoters and enhancers were enriched in different sets of binding sites (Figure 7B). The active promoter chromatin state was enriched in PPARG binding sites independent of the time point (Score = 37-55). These regions were enriched in CEBPB binding sites to a lesser extent (score = 16). Active and genic enhancer states were the most enriched in CEBPB binding sites (score = 18-30). Different enhancer states were enriched in cofactors EP300, MED1, and RXRG binding sites (score = 5-30). The enrichment of enhancers in these binding sites increased over time. These observations suggest that the cofactors may not share the same sites but bind to other regulatory regions of the same targets. This might explain the discrepancies between the factor-co-factor overlap of based on binding sites vs. gene targets.

Adipogenic transcription factors regulate autophagy through other transcription factors and kinases. PPARG targeted DNA-binding transcription factors, especially early on during the differentiation course (Figure-Figure 8A). We first observed that PPARG target several genes labeled as transcription factors and autophagy-related in the gene ontology terms. The effect was more significant (ratio = 0.3 at FDR *<* 0.2) in the case of autophagy compared to lipogenesis. CEBPB targeted a smaller number of these factors. Both factors targeted genes coding for protein kinases throughout the course of differentiation (Figure 8A). By contrast, the two factors, and CEBPB in particular, increasingly targeted genes coding for protein phosphatases related to lipogenesis.

**Figure 8:**
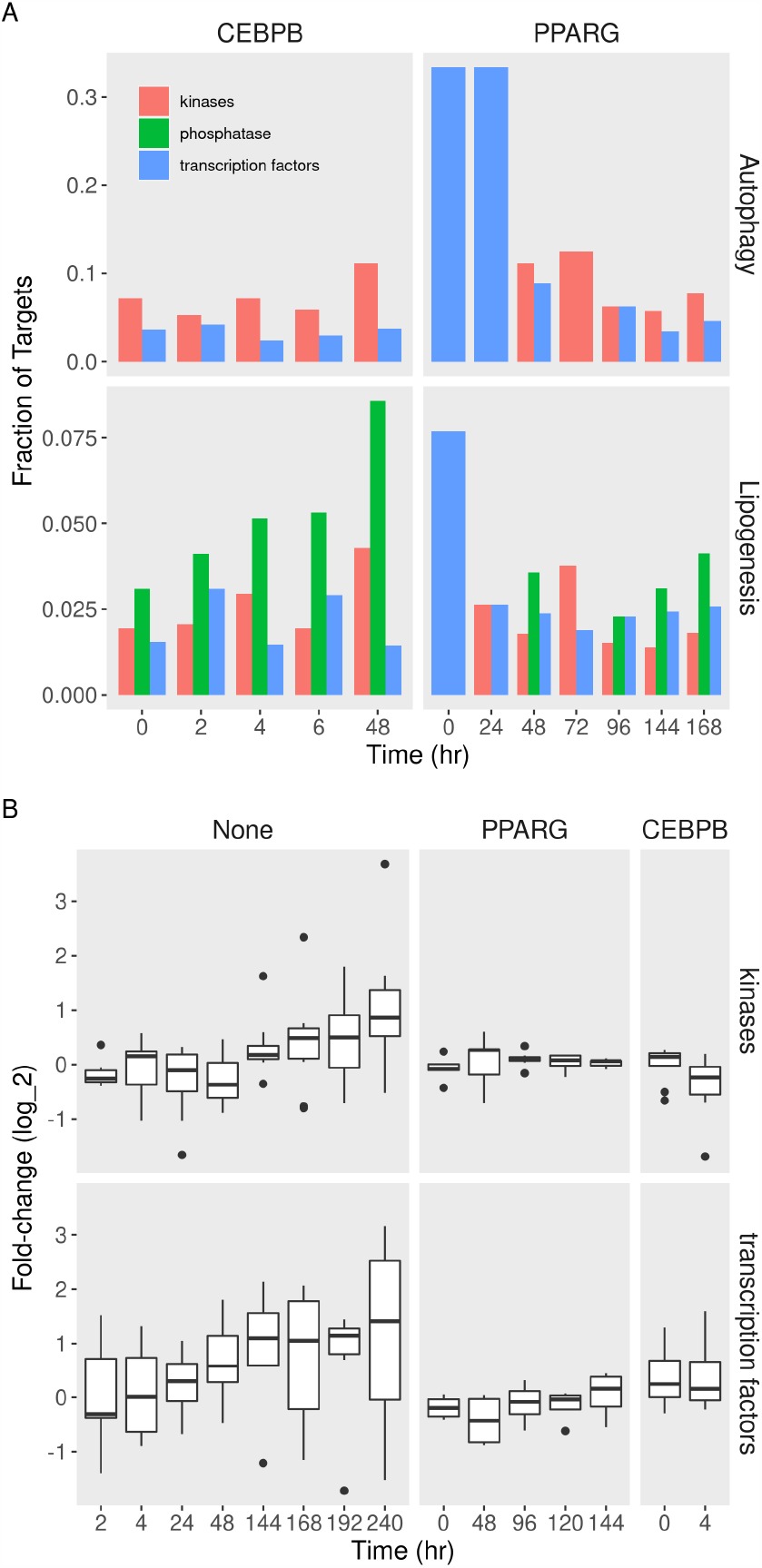
Fractions and expression of autophagy gene targets in different functions during adipogenesis. Binding peaks of PPARG or CEBPB were used to identify autophagy (and lipogenesis) gene targets at every time point of adipocyte differentiation. Read counts were used to quantify the gene expression at different times points of differentiation with or without perturbations. Expression was compared to pre-adipocytes (0 hr) to calculate fold-change and p-values. A) Numbers of gene targets in different molecular function terms (color). B) Fold-change (25, 50, and 75% quantiles) of the terms’ members in None, *Pparg*-or *Cebpb*-knockdown cells.

The expression of autophagy transcription factors gene and kinases was, on average, induced during the differentiation course (Figure 8B). The knockdown of *Pparg* in preadipocytes resulted in failed induction of this set of targets. This effect persisted for several days after the beginning of differentiation. A similar effect was observed for the knockdown of *Cebpb* at 4 hours.

## 4 Discussion

In this study, we used gene expression and chromatin binding data to build models for gene regulation and transcription factors sites/targets in differentiating adipocytes. We were able to identify likely targets among autophagy and lipogenic genes and evaluate the effect of their binding on expression. We also built multi-state models for the transcriptional regulators and chromatin states to explore the interactions between transcription factors, co-factors, and histone modifications. Figure 9 show diagrams of the main findings of the study.

**Figure 9:**
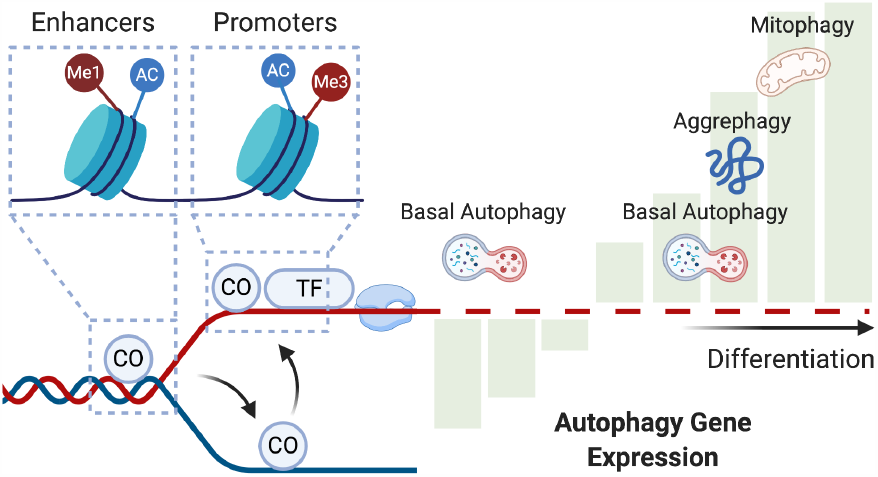
A model for transcriptional and chromatin modification on autophagy genes. Co-factors (CO) precede transcription factors (TF) to their shared targets. Co-regulators localize to enhancer regions marked by lysine monomethylation (Me1) and acetylation (Ac). They prime the targets for transcription, where transcription factors bind to the promoter regions marked by lysine tri-methylation (Me3) and acetylation (Ac). These events regulate the expression of autophagy genes in a bi-phasic manner. Early during adipogenesis, several autophagy genes are down-regulated, and possibly only basal autophagy is functional. Later, autophagy genes are up-regulated, and autophagy, organelle-specific autophagy, in particular, is activated.

We found that autophagy genes are regulated as part of the transcriptional program of differentiating adipocytes. Therefore, they might be regulated by the same adipogenic transcription factors (Figure 1). We previously made a similar observation [3]. Studies suggested several one-to-one links between those transcription factors. CEBPB induces the expression of *Pparg* either directly or by removing its inhibitors through autophagy [1]. We previously showed that adipogenic transcription factors CEBPB and PPARG regulate autophagy gene products during adipogenesis, either directly or indirectly through other transcription factors. Here, we further explore this regulation by examining the temporal and the spatial arrangement among those two factors, co-factors, and histone modifications.

Autophagy is essential for adipocyte differentiation. The knockdown of crucial autophagy genes such as *Atg5/7* resulted in failed induction of pre-adipocytes or reduced adipose mass tissue in mice [46, 47]. We observed the down-regulation of many autophagy gene products in the early days of adipogenesis (Figure 1). This is likely to impede many, but not all autophagy functions. Autophagy plays a role in maintenance and energy production in growing early adipocytes and possibly benefit the white adipocyte phenotype above others [48, 49]. In the later stages of differentiation, cells undergo phenotype changes that require the removal and recycling of intracellular components such as the mitochondria. Indeed, we observed the activation of organelle-specific forms of autophagy after two days of adipocyte induction (Figure 3). Together, the observed bi-phasic response to MDI-induction suggests two distinct autophagy functions in early and late-adipogenesis.

Although their expression increase in response to MDI, adipogenic factors have binding sites in pre-adipocytes (Figure 5B). CEBPB activates as early as 4 hours, and PPARG follows later during the differentiation course [50]. The abundance of the factors during the differentiation might explain this. The former is induced very early during adipogenesis, while PPARG levels do not max out until later. Co-factors exist on their targets irrespective of time points (Figure 6A). They might be able to access the majority of their targets at all times. The combination of factor-co-factor increased overtime for PPARG (Figure 6B). This is either because the complex is binding to more targets over time or because a combination of two proteins is necessary to induce the same targets.

Factors and co-factors localized to different genomic regions, even on the shared targets. As expected, transcription factors CEBPB and PPARG bind to the promoter regions the most. These regions were increasingly modified by histone markers associated with active promoters. PPARG could bind as a single factor suggesting a pioneering function [51]. In other words, it can access DNA, and other factors might provide selectivity. Co-factors were abundant in regions identified as promoters or enhancers (Figure 4). This suggests that co-factors such as RXRG and MED1 are required to bind with the main factors but may perform other roles. Those could be a form of assisted loading or priming enhancer regions for transcription factors binding [52].

A break down of the types of binding targets for the two adipogenic factors revealed an interesting pattern. PPARG targets the genes coding for other transcription factors with down-stream autophagy genes (Figure 8A). This was also evident in the case of lipogenic genes. CEBPB, on the other hand, was mostly bound to genes involved in other activities such as kinases and proteases. In addition to the larger number of targets, the high number of transcription factors of PPARG suggests a broader effect on regulating autophagy genes. The more downstream transcription factor genes, the larger the effect of the factor. Indeed, knocking down *Pparg* resulted in a broader range of dysregulation (Figure 8B).

Studies suggested that a specific arrangement of transcriptional regulators is required for successful reprogramming of differentiating neurons or neutrophils [53, 54]. Understanding the role of these regulators enables managing the differentiation process and the function of the differentiated cells. By manipulating certain factors, it would be possible to fine control the course of cell development. Inhibiting mitophagy in pre-adipocytes, for example, maintained the beige adipocyte phenotype, rather than the white, and resulted in cells with greater thermogenic capabilities [49]. Reversing the differentiation of adipocytes would also be possible either by targeting the regulators directly or the specific autophagy function they regulate [55]. Finally, mature adipocytes specialize in storing lipids, a function which lipophagy could modify.

Our analysis was limited to the time points for which data was available. For example, we observed that the CEBPB effectively targeted and affected the expression of autophagy genes as early as 4 hours. No data before 4 hours were available. Therefore we do not know whether this effect can be observed earlier. The curated datasets comprise data from previously published studies. We carefully curated and processed the data to reduce batch effects resulting from the variations among the studies. In the case of ChIP-seq data, we only used replicated peaks when more than one sample was available.*Pparg*-knockdown microarrays data had missing information on multiple genes that did not have corresponding probes. Finally, in the factor perturbation data during the differentiation course, it was difficult to disentangle the time from the perturbation effect.

## Acknowledgments

This study was supported by the National Research Foundation of Korea (NRF) grant funded by the Ministry of Science and ICT (MSIT) of the Korea government [2015R1A5A2008833 and 2020R1A2C2011416].

## Author contributions

MA conceived the idea of the study, curated the data, performed the analysis and wrote the first draft of the manuscript. THL, TMP, SZ, JSH, and OE contributed to the writing of the manuscript. DRK acquired the funding, supervised the project and contributed to the concepts of the study and the writing of the manuscript.

## Data availability

This study includes no data deposited in external repositories.

